# The epigenetic Oct4 gene regulatory network: stochastic analysis of different cellular reprogramming approaches

**DOI:** 10.1101/2023.03.01.530689

**Authors:** Simone Bruno, Domitilla Del Vecchio

**Affiliations:** Department of Mechanical Engineering, Massachusetts Institute of Technology, 77 Massachusetts Avenue, Cambridge, MA 02139. Emails: (,)

## Abstract

In the last decade, several experimental studies have shown how chromatin modifications (histone modifications and DNA methylation) and their effect on DNA compaction have a critical effect on cellular reprogramming, i.e., the conversion of differentiated cells to a pluripotent state. In this paper, we compare three reprogramming approaches that have been considered in the literature: (a) prefixed overexpression of transcription factors (TFs) alone (Oct4), (b) prefixed overexpression of Oct4 and DNA methylation “eraser” TET, and (c) prefixed overexpression of Oct4 and H3K9me3 eraser JMJD2. To this end, we develop a model of the pluritpotency gene regulatory network, that includes, for each gene, a circuit recently published encapsulating the main interactions among chromatin modifications and their effect on gene expression. We then conduct a computational study to evaluate, for each reprogramming approach, latency and variability. Our results show a faster and less stochastic reprogramming process when also eraser enzymes are overexpressed, consistent with previous experimental data. However, TET overexpression leads to a faster and more efficient reprogramming compared to JMJD2 overexpression when the recruitment of DNA methylation by H3K9me3 is weak and the MBD protein level is sufficiently low such that it does not hamper TET binding to methylated DNA. The model developed here provides a mechanistic understanding of the outcomes of former experimental studies and is also a tool for the development of optimized reprogramming approaches that combine TF overexpression with modifiers of chromatin state.

## 1 Introduction

Through the process of cellular differentiation, embryonic stem cells evolve into a variety of specialized cell types in multi-cellular organisms. The approach to convert differentiated cells to a pluripotent state (induced pluripotent stem cells, iPSCs) is called cellular reprogramming [1]. Since human iPSCs have functions almost identical to the ones of embryonic stem cells (ESCs), they can be used to replace damaged cells, representing a promising alternative to ESCs for regenerative medicine [2, 3]. Other applications of iPSCs include *in-vitro* disease modeling and drug screening/discovery [3]. The first iPSC reprogramming approach, introduced by Yamanaka and colleagues [4, 5], is based on prefixed overexpression of four key transcription factors (TFs), Oct4, Sox2, Klf4, and c-Myc (OSKM factors). Because the efficiency of the initial reprogramming process was very low (1-2 % [6–9]), a plethora of follow-up studies have appeared with an aim of improving efficiency [10–14].

We can largely group these studies into (a) those that keep the TF cocktail to OSKM and investigate the extent to which the levels of these factors influence efficiency [11, 15–20] and (b) those that add factors to the original OSKM cocktail, such as epigenetic modifiers [21–24]. In particular, the studies using epigenetic modifiers are grounded on the fact that in terminally differentiated cells, such as fibroblasts, used for cell reprogramming, the genes of the pluripoteny GRN are “shut off” often due to highly compactified chromatin [25–29]. In fact, in eukaryotic cells the DNA is wrapped around nucleosomes, octamers of proteins called histones. The extent to which DNA is more or less tightly wrapped around these nucleosomes determines how easily a gene can be transcribed [30]. Specifically, DNA structure (chromatin state) can be either more relaxed and transcriptionally-permissive (euchromatin) or more compressed and transcriptionally prohibitive (heterochromatin). The extent of chromatin compaction is dictated by specific enzymatic modifications to the histones and DNA, such as H3K9 methylation (H3K9me3) or H3K4 methylation/acetylation (H3K4me3/ac), and DNA methylation [25]. This implies that the chromatin state provides an additional layer of transcriptional regulation.

In the last decade, many experiments have been conducted to understand how chromatin modifications affect cellular reprogramming [21–24, 31, 32]. More precisely, these experimental studies aimed to determine which chromatin modifications have a key role on the reprogramming process and the best approaches to improve the process efficiency (percentage of reprogrammed cells at a fixed time) and reduce its stochasticity, or latency variability, that is, the variability of the “time that an individual cell takes until it gives rise to a daughter iPS cell”[33].

In this paper, we focus on TF Oct4, since it is well known that overexpression of Oct4 alone is sufficient for iPSC reprogramming [15, 19, 34–36] and that Oct4 is a key regulator of enzymes that remove DNA methylation (TET) and H3K9me3 (JMJD2) [25, 37–41]. We create a model for the three-gene network composed of Oct4, TET, and JMJD2 and define this network the epigenetic Oct4 gene regulatory network (Epi Oct4 GRN). We then analyze the model via simulation by using Gillespie’s Stochastic Simulation Algorithm (SSA) [42]. More precisely, in order to determine the efficacy of different reprogramming approaches, we analyze efficiency and latency variability. We also analyze how biochemical parameters, such as proliferation rate or the concentration of Methyl-CpG-binding domain proteins (MBDs), affect the stochastic behavior of the reprogramming process, with the aim of providing a mechanistic understanding of the outcomes that have been experimentally observed.

This paper is organized as follows. In Section 2, we introduce the Epi Oct4 GRN that we developed to conduct our study. In Section 3, we present the results of the computational analysis conducted. Finally, in Section 4 we present discussion and conclusive remarks.

### Related work

Some models that include histone modifications or DNA methylation into gene expression regulation to investigate iPSC reprogramming have appeared in the past years [43–46]. However, none of these models include both histone modifications and DNA methylation. Furthermore, none of these models include MBDs, whose presence has been shown to be critical for the effect of the reprogramming process [21, 31].

## 2 Model of the epigenetic Oct4 gene regulatory network

The traditional pluripotency GRN, responsible for the preservation of pluripotency, is made of three key TFs, Oct4, Sox2, and Nanog, which self-activate and mutually activate one another [47–49]. Here, we focus on Oct4 only and, in order to take into account epigenetic changes currently understood to take place during iPSC reprogramming, we include two more genes that express chromatin modifiers. These chromatin modifiers are JMJD2 and TET, enzymes involved in the erasure process of histone modifications H3K9me3 and DNA methylation [25, 37–41], two chromatin modifications associated with compacted chromatin state [50]. More precisely, while JMJD2 directly erases H3K9me3 [25, 37–39], TET enzyme recognizes CpGme dinucleotides and converts methylated CpG to carbolxylcytosine through multiple intermediate forms [40, 41], none of which is recognized by DNMT1, the enzyme responsible for copying the CpGme pattern on the nascent DNA strand during DNA replication [50]. Oct4 recruits writers of H3K4me3 to its own gene [51] and to the JMJD2 [52] and TET [53] genes. This leads to a model in which Oct4 self-activates and mutually activates TET and JMJD2 by recruiting writers of activating chromatin modifications, while TET and JMJD2 self-activate and mutually activate Oct4 by erasing repressive chromatin modifications. We call this GRN the epigenetic Oct4 gene regulatory network (Epi Oct4 GRN).

The chromatin modification circuit within each gene has been developed in [54]. This circuit includes H3K9 methylation (H3K9me3), DNA methylation (CpGme), H3K4 methylation/acetylation (H3K4me3/ac), and their known interactions. As mentioned above, the first two modifications are associated with a repressed gene state [50], while H3K4me3/ac is associated with an active gene state [25, 55]. Then, by considering this circuit for each gene, the expression rate of each gene will be determined by the number of nucleosomes with activating (D^A^) or repressive 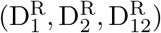 chromatin modifications (Fig. 1(b)).

**Figure 1:**
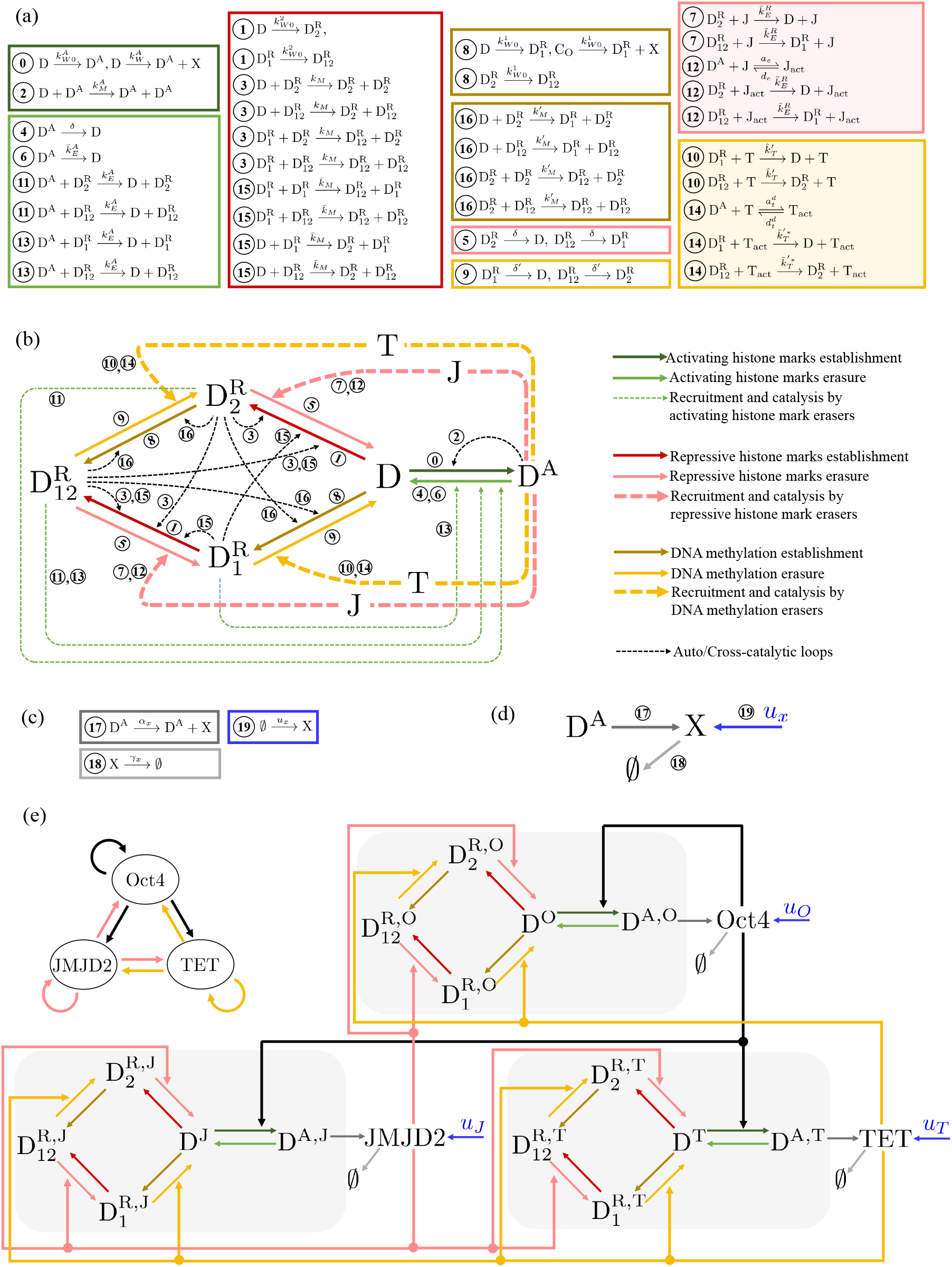
The epigenetic Oct4 gene regulatory network (Epi Oct4 GRN): circuit and reactions. (a) Reactions associated with the chromatin modification circuit of each gene X, with X = Oct4 (O), TET (T), JMJD2 (J). The reactions are described in Section 2. The reactions associated with activating histone modifications, repressive histone modifications and DNA methylation are enclosed in green boxes, pink boxes, and yellow boxes, respectively. Dark shades are associated with reactions describing the establishment of the modifications and light shades are associated with reactions describing the erasure of the modifications. Furthermore, shaded boxes enclose reactions involving T and J. (b) Diagram representing the chromatin modification circuit for each gene X, with X = O,T,J. (c) Reactions associated with the production (dark gray box), dilution/degradation (light gray box) and artificial overexpression *u_x_* (blue box) of the gene product X, with X = O,T,J. The numbers on the left hand side of the reactions are described in Section 2. (d) Diagram representing the production, dilution/degradation, and overexpression of X, with X = O,T,J. (e) Diagram of the Epi Oct4 GRN, in which Oct4 self-activates and mutually activates TET and JMJD2 by recruiting writers of activating chromatin modifications on all genes (black arrows), while TET and JMJD2 self-activate and mutually activate Oct4 by recruiting erasers for repressive chromatin modifications on all genes (pink and yellow arrows, respectively). Compared to panel (b), to simplify the diagram, we did not represent the dashed arrows indicating recruitment and catalysis in each gene’s chromatin modification circuit.

We next describe both the chromatin modification circuit and the model of gene expression (see [54] for a more detailed description). Both histone modifications and DNA methylation can be *de novo* established (process encapsulated in reactions 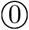, ①, and ⑧). Then, histone modifications can enhance the establishment of marks of the same kind to nearby nucleosomes via a read-write mechanism, generating auto-catalytic reactions (encapsulated in ②, ③). Analogously, repressive histone modifications enhance the establishment of DNA methylation, and viceversa, generating cross-catalytic reactions (encapsulated in 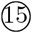, 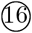). Finally, each modification can be passively removed through dilution, due to DNA replication (reactions ④, ⑤, and ⑨), or through the action of eraser enzymes (basal erasure) (reactions ⑥, ⑦, and ⑩). These erasers can be also recruited by the opposite modifications (recruited erasure), that is, repressive modifications recruit activating modification’s erasers and *viceversa* (reactions 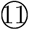, 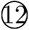, 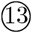 and 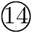). In this model the rate of the processes described above for H3K9me3 (DNA methylation) is assumed not to change if the other repressive mark is present on the same nucleosome. In terms of species, in this model we have D (nucleosome with DNA wrapped around it), D^A^ (nucleosome with H3K4me3/ac), 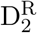 (nucleosome with H3K9me3), 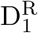 (nucleosome with CpGme), and 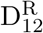, (nucleosome with both H3K9me3 and CpGme). Given that we are interested in wiring three chromatin modification circuits, each one of them associated with Oct4, TET, and JMJD2 genes, respectively, the other species included in our model are Oct4 (O), TET (T), and JMJD2 (J). All the reactions described above are collected in the list of Fig. 1(a), in which the reactions involving TET and JMJD2 are shaded in yellow and pink respectively. A diagram of the chromatin modification network considered is provided in Fig. 1(b). It is important to point out that in this system the transcriptional self-activation are modeled as a Hill function with cooperativity 1, that is 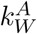 (Fig. 1(a)) is a monotonically increasing function of the abundance of X (X), that can be written as 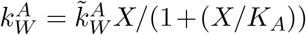, in which 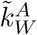 is a coefficient that does not depend on *X*, *K_A_* is the dissociation constant of the binding reaction between X and DNA, and Ω represents the reaction volume [54].

The transcriptional regulation model is also taken from [54]. Here, transcription is allowed only by D^A^, and transcription and translation are lumped together (reaction 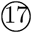). The trascription by D is considered negligible because it is assumed that transcription of D by RNA polymerase II occurs concurrently with H3K4me3 deposition (i.e., conversion of D to D^A^), as observed in [56]. Furthermore, the gene product X is subject to dilution due to cell division and degradation (reaction 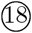). Finally, the production of X can be artificially increased through a prefixed overexpression (reaction 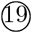). These reactions are listed in Fig. 1(c) and a representative diagram is shown in Fig. 1(d). Then, based on the interactions among Oct4, TET and JMJD2, we can wire the three chromatin modification circuits to obtain the Epi Oct4 GRN circuit that we analyze in this paper (Fig. 1(e)).

In order to better understand the analysis and the results presented in the next section, let us introduce the following parameters and variables. The first one is *D_tot_* = D_tot_/Ω, in which D_tot_ represents the total number of modifiable nucleosomes within the gene of interest and Ω represents the reaction volume, and the normalized time 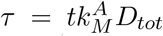. Let us define the dimensionless parameters 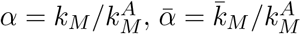, and 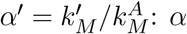 represents the normalized rate constant of the auto-catalytic reaction for repressive histone modifications, and 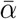 and α’ represent the normalized rate constants of the cross-catalytic reactions between repressive histone modifications and DNA methylation. Let us also introduce

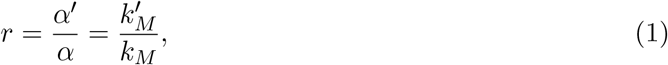

that is, the ratio between rates at which repressive histone modifications enhance the establishment of DNA methylation through cross-catalytic reactions and the rate at which repressive histone modifications enhance their own establishment through auto-catalytic reactions. Furthermore, we use *η* to represent the efficiency of the maintenance process of DNA methylation by DNMT1 [25], that is, *η* = *δ’/δ*, with *η* = 1 if DNMT1 is absent (no maintenance) and *η* = 0 if the maintenance process is 100% efficient. Now, we define

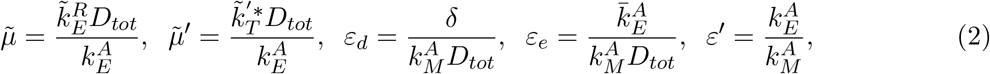

with 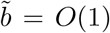 such that 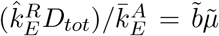 and 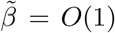 such that 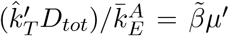. More precisely, 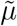 is a dimensionless parameter quantifying the asymmetry between the erasure rates of repressive and activating histone modifications, while 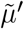 is a dimensionless parameter quantifying the asymmetry between the erasure rates of DNA methylation and activating histone modifications. Furthermore, 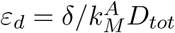 is the normalized rate constant associated with dilution due to DNA replication, and, since 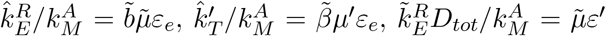 and 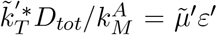, the dimensionless parameter *ε_e_* (*ε’*) scales the ratio between the rate of the basal erasure (recruited erasure) and the one of the auto/cross-catalytic reaction of each chromatin modification. Finally, related to the gene expression, we introduce 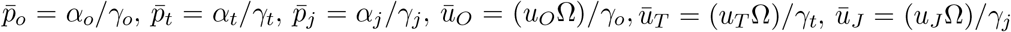 and 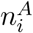, with i = O,T,J, that is, the total amount of nucleosomes modified with active chromatin modifications for each gene *i*. Finally, let us show the relationship between MBD proteins (B) and the rate coefficients of the reactions associated to the erasure of DNA methylation due to the action of TET. More precisely, these reaction rate coefficients are 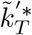 for the reactions in which TET is recruited by D^A^ (reactions 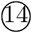) and 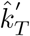 for the reactions in which TET is not recruited by D^A^ (reactions ⑩). As derived in [54], 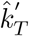 and 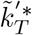 can be written as

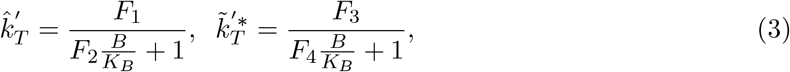

in which *K_B_* is the dissociation constant associated to the binding reaction between B and methylated DNA and *F*_1_, *F*_2_, *F*_3_ and *F*_4_ are functions that do not depend on B or *K_B_*. Then, 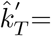 and 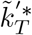 increase when *B/KB* decreases. Now, from (2), it is possible to notice that 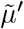 is an increasing functiong of 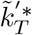. We can then write

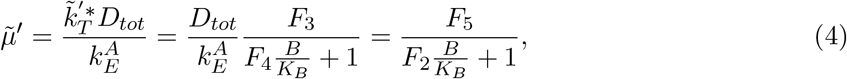

in which 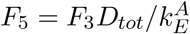 is a function that does not depend on B or *K_B_*. From (4), we can conclude that knocking down MBD proteins (*B* = 0) or locally preventing their binding to methylated DNA (*K_B_* → ∞) allow to increase 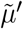.

## 3 Results

We first study how DNA methylation affects cellular differentiation, and then we compare different reprogramming approaches with respect to their efficiency and stochasticity. To this end, we perform a computational study of the temporal trajectories of the system by simulating the system of reactions associated with the Epi Oct4 GRN (Fig. 1(e)) with the Gillespie’s Stochastic Simulation Algorithm (SSA) [42].

### Effect of DNA methylation on differentiation

Here, we determine how the erasure rate constant of DNA methylation 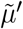 affects the first time that, without any external stimulus, the Oct4 gene reaches the repressed state 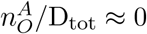 (corresponding to a differentiated state [35]), starting from the active state 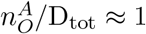 (as in the pluripotent state [35]) (Fig. 2). It is possible to notice that, for values of 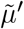 sufficiently large, none of the trajectories reach the Oct4 repressed state. By reducing 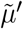, most of the trajectories reach, in a stochastic manner, the Oct4 repressed state after a finite time. Finally, if we keep reducing 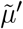, then the time trajectories reach 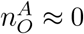 more quickly.

**Figure 2:**
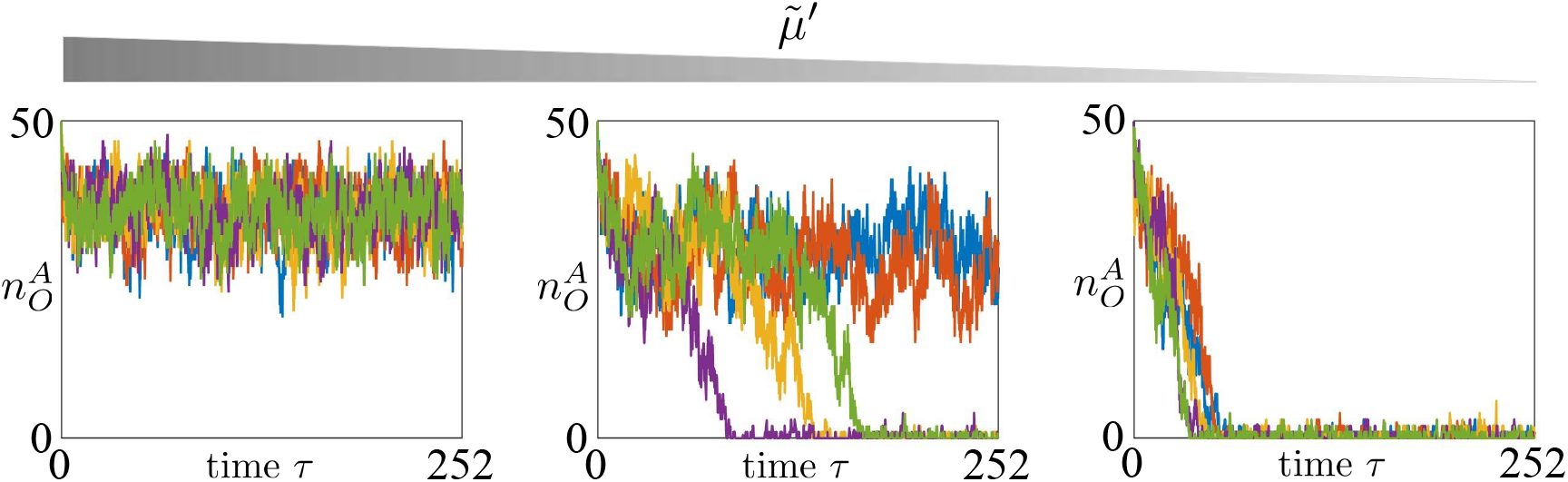
Small DNA methylation erasure rate leads to a faster differentiation process. Time trajectories of 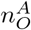 starting from the Oct4 fully active state for different values of 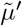. In all plots, on the *x* axis we have the time normalized with respect to 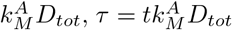. The parameter values used for these simulations can be found in SI-Table 1. In particular, we set 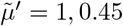, 0.2, *ε_d_* = 0.4, 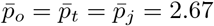, *η* = 0.2, *μ* =1, *ε_e_* = 0.4 and *ε’* = 1. In our model, parameter 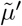 quantifies the asymmetry between the erasure rates of DNA methylation and activating histone modifications. Mathematical definition of 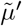 can be found in (2). For all the simulations shown in the figure, we considered D_tot_ = 50.

Overall, these results show that, if the DNA methylation erasure rate constant 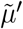 is sufficiently small, then the system will always reach the Oct4 gene repressed state. From a mathematical point of view, this implies that DNA methylation erasure rate sufficiently small makes the pluripotent state unstable. Even if previous models of the pluripotency GRN view the plutipotent state as a stable attractor of a multistable system [49, 57, 58], our results are in agreement with what, more in general, we observe in multicellular organisms, that is, pluripotent cells always differentiate after a certain finite time. This computational study also suggests that, in order to increase the efficiency of a reprogramming process, one potential approach is to increase the rate at which DNA methylation is erased (making 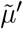 higher), since this improves the stability of the Oct4 active gene state (Fig. 2).

Finally, given that H3K9me3 decay rate is estimated to be much higher than the decay rate of DNA methylation [54, 59], decreasing the erasure rate constant of repressive histone modifications 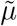 does not affect the speed of the differentiation process as much as decreasing DNA methylation erasure rate (See SI-Fig. S.1).

### Effect of proliferation rate on reprogramming through Oct4 overexpression

Here, we analyze how the proliferation rate *ε¿* affects the efficiency of the reprogramming process based on Oct4 overexpression. To this end, we analyze the trajectory of the active chromatin state of the Oct4 gene, 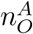, starting from a fully repressed state 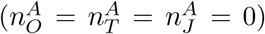, when we artificially overexpress Oct4, that is, we set 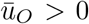 in the reactions listed and represented in Fig. 1(c),(e). We then define the process efficiency as %PL, that is the percentage of time trajectories of 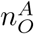 that reach the active state by a fixed time point and we evaluate, for a fixed input 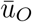, %PL for different values of *ε_d_*. The analysis shows that increasing *ε_d_* speeds up the process and this prediction is in agreement with experiments conducted in [33], showing that a higher ploriferation rate makes the reprogramming process faster. This is because increasing *ε_d_* leads to higher decay rate of all the modifications (Equation (2) and Fig.1). Given that the initial state is the Oct4 repressed state, mainly characterized by repressive chromatin marks, then higher *ε_d_* leads to a faster erasure of DNA methylation and repressive histone modifications. This, in turn, allows activating histone modifications to establish faster and leads to a faster reactivation process.

### Reprogramming based on concurrent Oct4 and TET overexpression

Now, we consider a reprogramming approach in which the enzyme TET is also overexpressed. We first determine how efficiency %PL and the latency variability are affected by different levels of TET overexpression 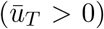. To this end, we conduct several simulations for a fixed 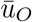 (fixed level of Oct4 overexpression) and different values of 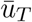 (Fig. 4(a)). The simulations show that adding overexpression of TET makes the Oct4 reactivation process faster and reduces the latency variability. This is because activating histone modifications can be established only on unmodified nucleosomes, as shown in the chromatin modification circuit diagram (Fig. 2(b)), and therefore the reprogramming process cannot start until repressive modifications are erased. Furthermore, while Oct4 is the protein recruiting the writer enzymes for the active chromatin modifications, TET itself is an erasure enzyme for DNA methylation. Therefore, overexpressing TET boosts the erasure of DNA methylation, accelerating the overall process.

**Figure 3:**
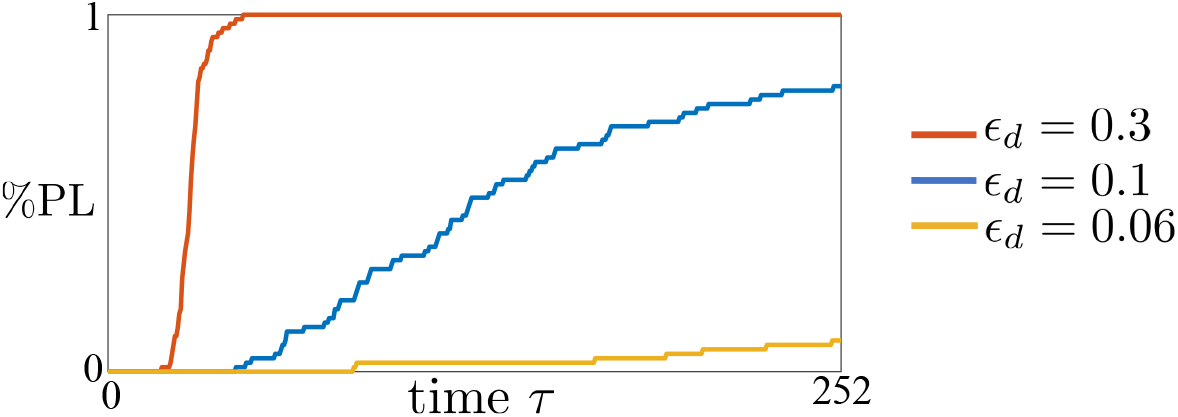
Higher proliferation rate speeds up the reprogramming process through Oct4 overexpression 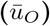. %PL for different values of *ε_d_*. The parameter values used for these simulations can be found in SI-Table 1. In particular, we consider *ε_d_* = 0.3, 0.1, 0.06 and we set 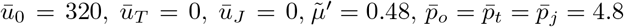, *η* = 0.2, *ε_e_* = 0.4, 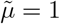 and *ε’* = 1. Here, we use lower values of *ε_d_* compared to the one used in Fig. 2 in order to take into account that proliferation decreases during differentiation [60]. Same choice will be done for the simulations shown in all next figures. In our model, parameter *ε_d_* represents the normalized rate constant associated with dilution due to DNA replication. Mathematical definition of *ε_d_* can be found in (2). For all the simulations shown in the figure, we considered a time span of 21 days (*τ* = 252) and D_tot_ = 50.

**Figure 4:**
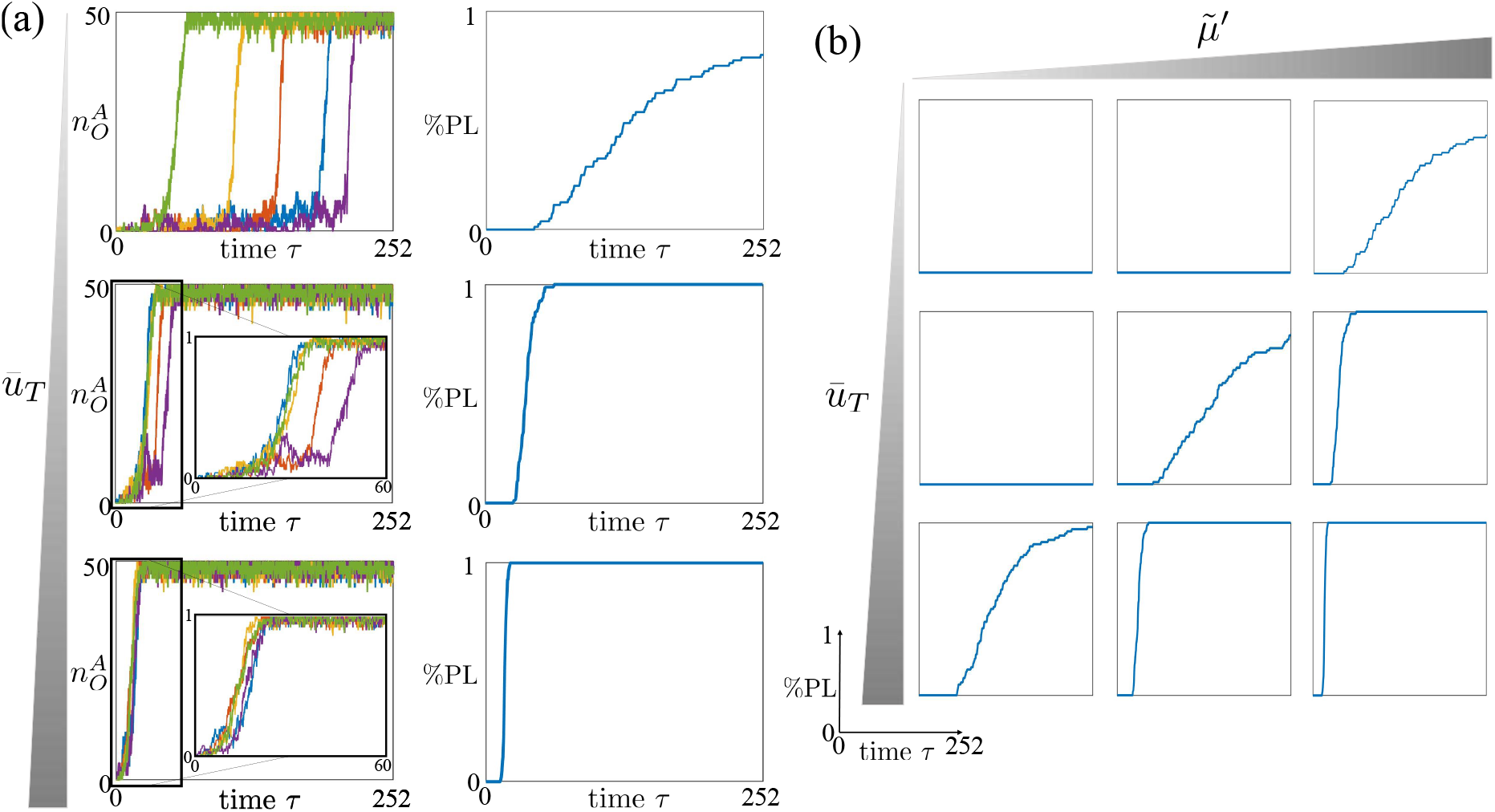
Concurrent Oct4 and TET overexpression lead to a more efficiency and less stochastic reprogramming process, under specific parameter regimes. (a) Left hand side plots: time trajectories of 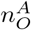 starting from the Oct4 fully repressed state 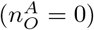 for different values of 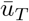. Right hand side plots: %PL, that is, the normalized amount of *N* = 100 time trajectories which reach 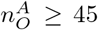, starting from 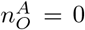. In all plots, on the *x* axis we have the time normalized with respect to 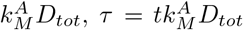. The parameter values used for these simulations can be found in SI-Table 1. In particular, we consider three values of 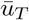 (i.e., 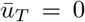), and we set 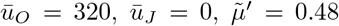, *ε_d_* = 0.1, *ε_e_* = 0.4, 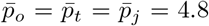, *η* = 0.2, 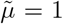 and *ε’* = 1. (b) %PL for different values of 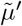 and 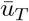. The parameter values used for these simulations can be found in SI-Table 1. In particular, we consider 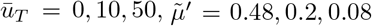, and all the other parameter values equal to the ones considered for the simulations in panel (a). In our model, 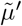 quantifies the asymmetry between the erasure rates of DNA methylation and activating histone modifications. Mathematical definition of 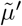 can be found in (2). For all the simulations shown in the figure, we considered a time span of 21 days (*τ* = 252) and D_tot_ = 50.

However, experimental data showed that adding TET overexpression to TFs overexpression does not increase iPSC reprogramming efficiency so pronouncedly [24]. Based on experimental studies previously conducted [21, 31] and according to the chromatin modification circuit [54] that we used in our Epi Oct4 GRN, one possible cause is the presence of MBD proteins. In fact, since MBD proteins bind to methylated CpG dinucleotides [61] and then protect them from being bound by TET [62], TET overexpression does not enhance the erasure process of DNA methylation unless MBD proteins are prevented from binding DNA. In fact, knocking down MBD proteins or locally preventing their binding to methylated DNA corresponds to a higher DNA demethylation rate constant 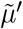 (See Section 2, Eq. (2)). In order to verify how different levels of MBD affect the TET overexpression reprogramming process, we evaluate the %PL of the Oct4 gene reactivation process for several values of 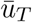 and 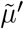 (Fig. 4(b)). The higher 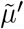 is (obtained by lowering the MBD proteins level), the faster and less stochastic the reactivation process is. These results are consistent with previous experimental data showing that global knowck down of MBDs increases significantly iPSC reprogramming efficiency [21, 31]. Furthermore, these results also show that with high level of MBD proteins (low 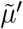), the iPSC reprogramming is slower and more stochastic, but for a sufficiently high TET overexpression level (high input 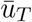), the process may still have an almost constant latency (Fig. 4(b)). However, it could be possible that these high values of TET overexpression cannot be achieved in practice.

### Reprogramming based on concurrent Oct4 and JMJD2 overexpression

From the previous analysis, we observe that, compared to overexpression of Oct4 alone, the addition of TET overexpression, that is responsible for the erasure of DNA methylation, leads to iPSC reprogramming faster and to decreased stochasticity of the process. Based on experimental studies conducted in the last decade, the repressive histone modification H3K9me3 seems to be a similarly crucial barrier for the reprogramming process [22, 63].

Then, we evaluate how the process efficiency and latency variability vary for three different levels of JMJD2 overexpression, that is, three different values of input 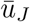 (Fig. 5(a)). Furthermore, in order to properly compare this reprogramming approach with the one based on the concurrent Oct4 and TET overexpression, for the parameter 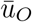 (Oct4 overexpression level) we consider the same value used in the previous analysis. The simulations show that JMJD2 overexpression makes the reprogramming process faster and reduces the latency variability (Fig. 5(a)). These results are consistent with experimental data showing how the addition of JMJD2 to the OSKM cocktail increases the iPSC reprogramming efficiency [22].

**Figure 5:**
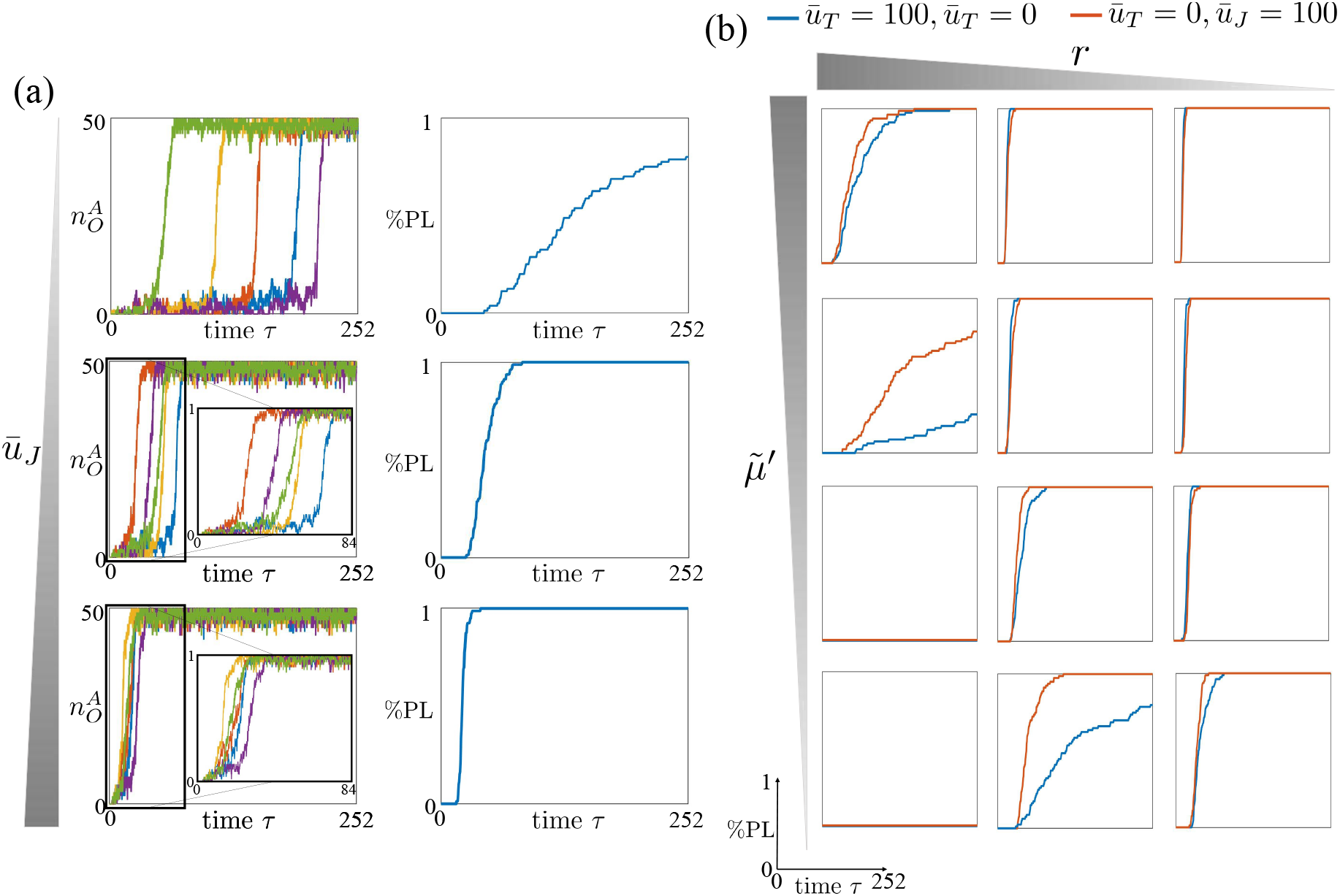
JMJD2 and TET overexpression can have different effect on the reprogramming efficiency and stochasticity. (a) Left hand side plots: time trajectories of 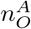 starting from the Oct4 fully repressed state 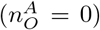 for different values of 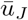. Right hand side plots: %PL, that is, the normalized amount of *N* = 100 time trajectories which reach 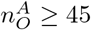, starting from 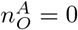. In all plots, on the *x* axis we have the time normalized with respect to 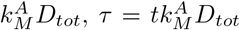. The parameter values used for these simulations can be found in SI-Table 2. In particular, we consider three values of 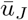 (i.e., 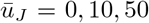), and we set 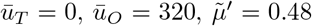, *ε_d_* = 0.1, *ε_e_* = 0.4, 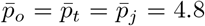, *η* = 0.2, 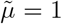 and *ε’* = 1. (b) %PL for different values of 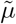 and *r*. The parameter values used for these simulations can be found in Table 2. In particular, we consider 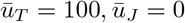 (blue lines), 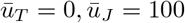 (red lines) and, for both cases 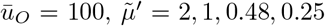, *r* = 5,1, 0.5, 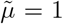, *ε_e_* = 0.4, 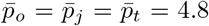, *η* = 0.2 and *ε’* = 1. In our model, 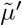 quantifies the asymmetry between the erasure rates of DNA methylation and activating histone modifications and *r* the ratio between rate at which repressive histone modifications enhance the establishment of DNA methylation through cross-catalytic reactions and the rate at which repressive histone modifications enhance their own establishment through auto-catalytic reactions. Mathematical definitions of *r* and 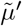 can be found in (1) and (2), respectively. For all the simulations shown in the figure, we considered a time span of 21 days (*τ* = 252) and D_tot_ = 50.

We next conduct a study to compare the latency of the Oct4 - TET overexpression approach to that of the Oct4 - JMJD2 overexpression approach (Fig. 5(b)). The results show how the rate and the stochasticity of the two reprogramming processes are affected by 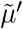 and *r* = *α’/α*. The parameter *r* is the ratio between the rate of the auto-catalytic reaction with which repressive histone modifications enhance their own establishment and the rate of the cross-catalytic reaction with which repressive histone modifications enhance the establishment of DNA methylation (See Equation 1).

In particular, for high values of 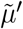, the addition of JMJD2 overexpression, compared to TET overexpression, leads to a faster reactivation of the Oct4 gene when *r* ≫ 1 (that is, repressive histone modifications enhance more the establishment of DNA methylation than their own establishment). This is because, if 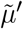 and *r* are both sufficiently high, then overexpression of JMJD2 leads to a fast erasure of repressive histone modifications. Then, without repressive histone modifications and their strong enhancement of DNA methylation, JMJD2 overexpression leads also to a fast erasure of DNA methylation. In this case, a reprogramming approach based on Oct4 - JMJD2 overexpression could be more efficient and have a less variable latency compared to the one based on Oct4 - TET overexpression (Fig. 5(b)).

By reducing 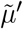, the Oct4 - JMJD2 overexpression approach becomes more efficient even for lower values of *r*. This is because, when DNA methylation enhancement by H3K9me3 is non-negligible (*r* sufficiently high), then the overexpression of TET may not be strong enough to erase DNA methylation if 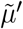 is low. On the other hand, for the same overexpression level, JMJD2 overexpression can be sufficient to erase repressive histone modifications and, together with them, their non-negligible enhancement of DNA methylation establishment (i.e., values of *r* sufficiently high) (Fig. 5(b)).

Overall, these results suggest that, due to the enhancement of DNA methylation by the positive reinforcement with H3K9me3, the addition of JMJD2 to Oct4 overexpression may be more effective than the addition of TET. However, for 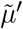 sufficiently high, Oct4 - TET overexpression approach is always more efficient than the Oct4 - JMJD2 overexpression approach.

## 4 Discussion

In this work, we first introduced the epigenetic Oct4 gene regulatory network (Epi Oct4 GRN), a network comprising a unique TF gene, Oct4, and two more genes expressing chromatin modifiers. These three genes (Oct4, TET, and JMJD2) are positively autoregulated and mutually activate each other, although through different mechanisms. Specifically, Oct4 self-activates and mutually activates TET and JMJD2 by recruiting writers of activating chromatin modifications, while TET and JMJD2 self-activate and mutually activate Oct4 by erasing repressive chromatin modifications. For each gene, we considered a chromatin modification circuit previously developed [54] (Section 2 and Fig. 1(b) and (c)). We then conducted a computational analysis to study three reprogramming approaches based on the overexpression of Oct4 alone, overexpression of Oct4 and TET enzyme, and overexpression of Oct4 and JMJD2 enzyme, respectively, by using Gillespie’s Stochastic Simulation Algorithm (SSA) [42] (Section 3).

Our analysis suggests that, for the same Oct4 overexpression level, the reactivation of the Oct4 gene is slower and more stochastic when only Oct4 is overexpressed (Fig. 3(a)) compared to the cases in which TET enzyme and JMJD2 enzyme are also overexpressed (Fig. 4(a) and Fig. 5(a)). Comparing the two latter cases, our results suggest that the former is more efficient if the recruitment of DNA methylation by H3K9me3 is sufficiently weak (*r* sufficiently low) and DNA methylation erasure is sufficiently fast (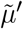 sufficiently large) (Fig. 5(b)). Experiments will be required to estimate the parameter *r* and determine which one is the more plausible scenario.

We also conducted a computational study to determine how the dosage of Oct4 overexpression affects the reprogramming process. The obtained results suggest that higher levels of Oct4 lead to a faster and more efficient reprogramming (See SI-Section S.3). However, previous studies based on the traditional pluripotency GRN models, including three TFs, Oct4, Sox2, and Nanog, have shown that a specific, intermediate Oct4 overexpression level may be needed for the success of the reprogramming process [49, 64]. This is achieved in models with three or more stable steady states, in which the pluripotent state is not extremal [49]. This is also consistent with experimental studies showing that an intermediate Oct4 level is required for pluripotency maintenance [65]. Our model, including only the Oct4 gene, is monostable, or at most bistable [54], in which the stable steady state can either be the somatic or pluripotent fate. Therefore, although this model predicts that increased Oct4 overexpression will lead to faster and more efficient reprogramming, this may not be the case in a tri-stable Oct4-Nanog-Sox2 model. Future studies will thus need to combine the Oct4-Nanog-Sox2 multistable model, with the chromatin modification circuit to enable concurrent investigation of the effect of dosage and chromatin state.

The contribution provided by this study not only allows us to compare different reprogramming approaches, but also provides a mechanistic understanding of multiple experiments conducted in the past years. The Epi Oct4 GRN model developed in this paper and the analysis conducted may thus aid the rational design of new gene reactivation approaches and their application to cell fate reprogramming.

## Acknowledgments

This work was supported by the National Institutes of Health (NIH/NIBIB Grant Number R01EB024591).

